# Sensory-motor computations in a cortico-pontine pathway

**DOI:** 10.1101/2022.02.22.481514

**Authors:** Timothy R. Darlington, Stephen G. Lisberger

## Abstract

Subcortical circuits decode the output from multiple cortical areas to create commands for movement. By recording during monkeys’ smooth pursuit eye movements, we reveal how the dorsolateral pontine nucleus (DLPN) and nucleus reticularis tegmenti pontis (NRTP) decode cortical output. Preparatory activity is inherited from the smooth eye movement region of the frontal eye fields (FEF_SEM_). Consistent with its effects on eye movement responses to visual motion, preparatory activity modulates the gain of pontine responses to visual motion input from visual area MT. Furthermore, the “place” code for target speed in area MT becomes the motor command’s “rate” code in the pons, through linear weighting of MT’s output. The division into decoding that happens in the pons and residual computations that occur downstream reveals that the pons is neither a relay nor the source of a final motor command. It is part of a complex, distributed cortical decoder for motor control.

## Introduction

The cerebral cortex is the source of many neural signals that influence the control of movement, including signals that encode expectation (Darlington et al., 2018), reward (Joshua & Lisberger, 2012; Kubanek & Snyder, 2015; Platt & Glimcher, 1999; Raghavan & Joshua, 2017), attention (Cohen & Maunsell, 2011), and sensory information (Omrani et al., 2016). Signals from disparate cortical areas must be combined to generate a motor command. Furthermore, the coordinate system of the cortical representation often requires a transformation to align with the coordinate system of the muscles that control movement (Groh & Sparks, 1992). In some systems, motor-related activity in the cortex has been understood as a low-dimensional dynamical system (Churchland et al., 2012; Churchland et al., 2010). However, it remains unclear how downstream brainstem and spinal circuits decode the output of a dynamical system to generate appropriate movements. Thus, a major challenge is to understand how sub-cortical neural circuits transform distributed cortical representations to create the appropriate commands for movement when it occurs. Here, we reveal how cortical signals are transformed as they pass through a cortico-pontine pathway to generate accurate motor output.

We evaluate the transformation of cortical population representations into commands for movement using smooth pursuit eye movements, an ideal model sensory-motor system for many reasons (Lisberger, 2010). We know much about the anatomical architecture of both the cortical and sub-cortical pathways for pursuit. We possess quantitative hypotheses about the functions of parallel cortical output pathways and how they contribute to the ultimate movement. We can characterize the motor output quantitatively and therefore draw rigorous conclusions about how subcortical pursuit areas combine multiple parallel outputs from the cerebral cortex to generate an appropriate eye movement response.

At least two parallel cortico-pontine output pathways play different roles in generating motor commands for pursuit. The middle temporal area (MT) is the main source of visual motion input to the pursuit system and sends projections, both directly and through area MST, to the pons (Distler et al., 2002; May & Andersen, 1986). Neurons in MT respond selectively to visual motion and are tuned for its speed and direction (Newsome et al., 1988), electrical micro-stimulation of area MT affects smooth eye velocity (Groh et al., 1997), and inactivation of area MT causes large deficits in smooth pursuit (Dürsteler & Wurtz, 1988; Dürsteler et al., 1987; Newsome et al., 1985). The smooth eye movement region of the frontal eye fields (FEF_SEM_) controls the strength, or “gain”, of visual-motor transmission and regulates access of visual motion signals to the oculomotor machinery. Sub-threshold stimulation in FEF_SEM_ enhances the behavioral responses to brief sinusoidal perturbations of target motion during fixation, when the perturbations normally are only weakly effective (Nuding et al., 2009; Tanaka & Lisberger, 2001). The firing of neurons in FEF_SEM_ is monotonically related to eye velocity and inactivation leads to large deficits in pursuit initiation and maintenance (Keating, 1991; MacAvoy et al., 1991; Shi et al., 1998).

Visual-motor gain and FEF_SEM_ have been implicated in many higher-order features of pursuit, for example reward (Raghavan & Joshua, 2017), motor learning (Li & Lisberger, 2011), and modulation of behavior by experience-dependent expectation of target motion (Darlington et al., 2018). Importantly, both visual-motor gain and preparatory activity in FEF_SEM_ are dialed up in preparation for smooth pursuit eye movements (Darlington & Lisberger, 2020; Tabata et al., 2006). It is possible to probe visual-motor gain by measuring the eye movement responses to brief pulses of target motion. These reveal an enhancement of gain that correlates well across time with the evolution of preparatory activity in FEF_SEM_, including larger eye movement responses when preparatory activity is larger under expectation of faster target speed.

We now attempt to move from a conceptual understanding of visual motion estimation and gain control to an understanding of how the brain combines these two signals to ultimately generate a motor command. Both MT and FEF_SEM_ project to the pons. At least anatomically, the pons receives a generous amount of cortical convergence suggesting that it might play some role in integrating visual motion and visual-motor gain signals (Brodal, 1980a; Distler et al., 2002; May & Andersen, 1986). The nucleus reticularis tegmenti pontis (NRTP) and the dorsolateral pontine nucleus (DLPN) are two pontine areas that have been linked to smooth pursuit eye movement. Lesions of either area cause deficits in smooth pursuit eye movements, and electrical micro-stimulation in either area can initiate smooth pursuit (May et al., 1988; Suzuki et al., 1999; Yamada et al., 1996). Previous neural recordings in NRTP and DLPN were limited by the use of only a very basic pursuit task or very basic passive visual motion stimuli without manipulating motor preparation (Mustari et al., 1988; Mustari et al., 2009; Ono et al., 2005; Suzuki et al., 2003). Thus, it is necessary to determine how cortical inputs are processed in the pons, especially to distinguish between visual motion and visual-motor gain related signals and understand how they interact.

In the present paper we revisit pursuit-related responses in NRTP and DLPN with a new set of stimulus and behavior paradigms to gain new insight into what roles they play in visual motion and visual-motor integration. We elucidate which parts of the transformation from cortex to pursuit occur in or before these pontine nuclei, and which must be handled by downstream circuits. Our analysis unfolds in 3 steps. First, we demonstrate sufficiency of these two nuclei by showing that the output of both is appropriate to drive the initiation of pursuit across a wide range of stimulus conditions. Second, we discover preparatory activity in both DLPN and NRTP that seems to be inherited from FEF_SEM_, and we show that the preparatory activity modulates the size of visual responses in the pons, in line with the effects on eye velocity. Third, we reveal that activity in the pons reflects an essentially complete transformation from the “place code” for target speed in MT to the rate code needed for pursuit, across target speeds and contrasts, and that this transformation can be performed by a surprisingly simple weighted sum of MT outputs. The division into transformations that occur before versus after the pons is striking and unexpected and reveals critical details of how the brainstem decodes cortical population responses to drive movement.

## Results

### NRTP and DLPN responses during pursuit initiation are consistent with an eye velocity motor command

Across a wide range of target speeds and two stimulus contrasts, the responses of NRTP and DLPN neurons during pursuit initiation closely track the eye velocity motor output. Visual comparison of the average population PSTHs (Figures 1C, D, F, and G) and the eye velocity (Figures 1I and J) show a very close qualitative match between the dynamics and amplitude of neural responses (Figure 1C, D, F, G) and behavioral responses (Figure 1I, J), across a speed range of 2, 4, 8, 16, and 32 deg/s, and for both high- and low-contrast targets. The responses to 32 deg/s are particularly revealing. For example, the behavioral latency and eye acceleration was longer and slower for target motion at 32 deg/s compared to 16 deg/s (Figure 1I and J, compare purple and red curves). The neural responses in both NRTP and DLPN mirror the latency and acceleration differences between 32 and 16 deg/s (Figures 1C, D, F, and G, compare purple and red curves). The population PSTH response latencies lead the behavioral response latencies slightly for NRTP and by a considerable amount for DLPN (downward arrows in Figure 1I, J and vertical dashed lines in Figure 1C, D, F, G).

**Figure 1.**
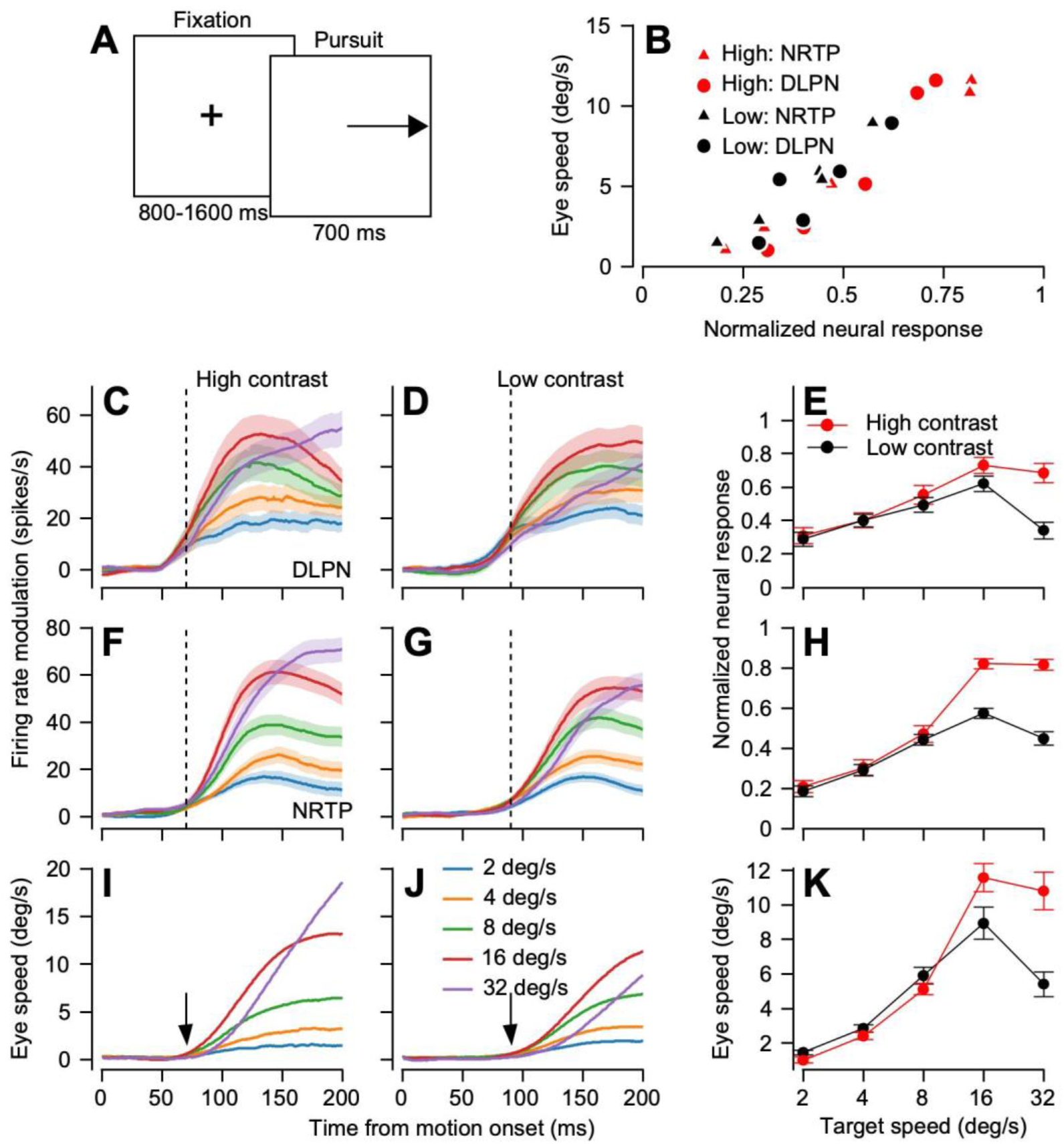
Responses of DLPN and NRTP neurons scale with eye velocity in pursuit initiation as function of target speed and contrast. **A**: Schematic of target motion for an individual pursuit trial. **B**: Red and black symbols plot normalized neural responses of NRTP (triangles) and DLPN (circles) neurons as a function of eye speed to high- and low-contrast targets, respectively. **C:** Different color traces plot the population-averaged firing rate of DLPN neurons during pursuit initiation in response to high-contrast targets moving at different speeds. **D**: Same as C for low-contrast targets. **E**: Red and black symbols connected by lines plot speed-tuning of normalized firing rate averaged across DLPN neurons to high- and low-contrast targets. **F**: Same as **C** for NRTP. **G**: Same as **G** for NRTP. **H**: Same as **E** for NRTP. **I:** Same as **C**, for eye velocity during pursuit initiation for high-contrast targets. **J:** Same as **I** for low-contrast targets. **K**: Red and black symbols connected by lines plot speed tuning of eye speed averaged at the end of pursuit initiation to high and low-contrast targets. Error-bands and error-bars represent the S.E.M.

Quantitative analysis shows how tightly the neural response amplitudes tracked the behavioral responses. For target speeds of 2, 4, and 8 deg/s, contrast did not affect either eye velocity (Figure 1K) or firing rate (Figure 1E, H) and both increased as a function of speed. Low-contrast targets caused smaller eye velocity and firing rates at 16 deg/s. Increasing target speed to 32 deg/s caused both firing rate and eye velocity to plateau for high-contrast targets and decrease for low-contrast targets. Finally, replotting the eye speed and neural response magnitudes across target speeds and contrast from Figures 1E, H, and K demonstrates a tight correlation that is independent of contrast and very similar for DLPN and NRTP (Figure 1B; NRTP: Pearson’s correlation coefficient = 0.98, p = 2.72 × 10^−7^, n = 10; DLPN: Pearson’s correlation coefficient = 0.93, p = 8.99 × 10^−5^, n = 10). We conclude that the output of NRTP and DLPN during pursuit forms an early eye velocity motor command.

### Motor preparatory signals propagate from FEF_SEM_ to pursuit-related pontine areas

We now turn our attention to understanding how visual motion and visual-motor gain signals are combined to form this motor command. Visual motion signals don’t have unfettered access to the oculomotor machinery, and pursuit responses to visual motion are subject to modulation by the speed of ongoing pursuit, expectations of target speed, reward, and other factors. Motor preparation and expectation of target speed are represented in the activity of neurons in FEF_SEM_ (Darlington et al., 2018; Mahaffy & Krauzlis, 2011). We now demonstrate that the preparatory activity in FEF_SEM_ propagates to both DLPN and NRTP.

Our experimental strategy is designed to separate the processing of visual information from its modulation by visual-motor gain. This separation is not possible during the initiation of pursuit, when both gain and visual input are changing dynamically. Therefore, we turn to a strategy of intentionally modulating the level of sensory-motor gain with preparation for movement and control of target speed expectation. We probe gain control without changing the setting of gain by delivering brief pulses of target motion under different conditions of motor preparation. We evaluate processing of visual information when gain is static, with passive visual motion during fixation.

We showed previously that the magnitude of eye speed during pursuit initiation can be explained as the result of a reliability-weighted combination of sensory input with expectations based on the history of target speed. We did so with a speed context paradigm (Figure 2B), where target motions were presented in blocks of trials that had either mostly fast (20 deg/s) or mostly slow (2 deg/s) target motions. Each target motion was preceded by a fixation interval of a duration randomized between 800 and 1600 ms (Figure 2A). During fixation, FEF_SEM_ neurons exhibit preparatory-related ramps of firing rate (Figure 2C). The preparatory activity is heterogeneous in the sense that it could be either an increase or a decrease in firing (Figure 2C). In addition, the ramps encode the expectation of target speed. On a neuron-by-neuron basis, the modulation of preparatory activity is greater during the fast-speed context compared to the slow-speed context (Figures 2F and I; paired sample t-test, p = 2.49 × 10^−20^, n = 322, Cohen’s d = 0.55).

**Figure 2.**
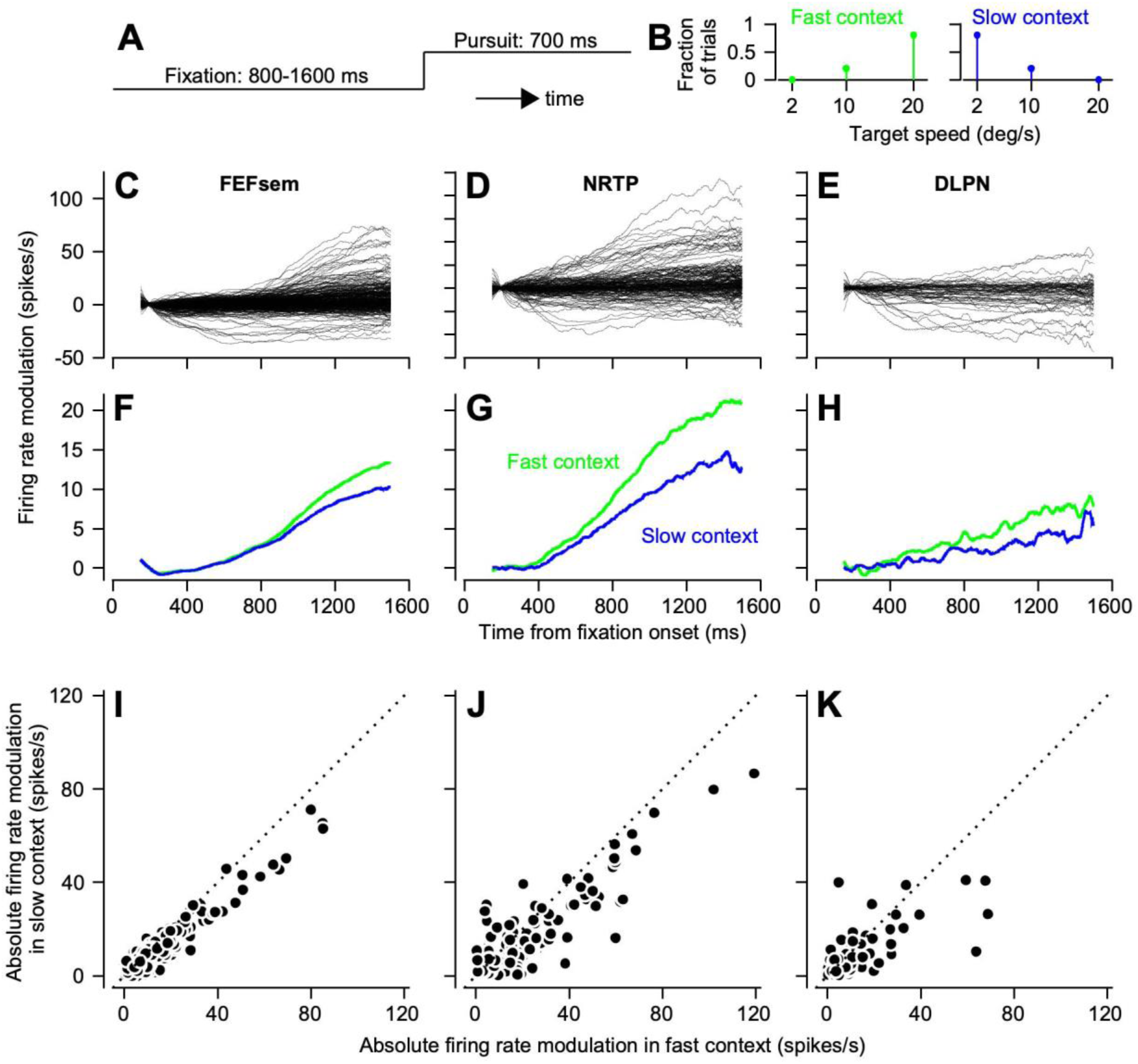
Preparatory activity is present in NRTP and DLPN during steady fixation. **A:** The timing of fixation and pursuit within a trial. **B**: The distribution of target speeds encountered in the fast- and slow-contexts. **C**: Traces plot the trial-averaged preparatory firing rate of individual FEF_SEM_ neurons as a function of time during fixation **D**: Same as **C** for NRTP. **E**: Same as **C** for DLPN. **F**: Red and black curves plot the firing rate averaged over all FEF_SEM_ neurons that exhibited preparatory increases of firing rate as a function of fixation time in the fast- and slow-contexts. **G**: Same as **F** for NRTP. **H**: Same as **F** for DLPN. **I:** Symbols plot the firing rate modulation across fixation during the fast context compared to the slow context for each neuron in our sample from FEF_SEM._ The dotted line plots the unity line. **J**: Same as **I** for NRTP. **K**: Same as **I** for DLPN. Panels C, F, and I are adapted with permission from Darlington et al., 2018.

Neurons in NRTP and DLPN display a heterogenous mix of increasing and decreasing firing rates during fixations leading up to pursuit (Figures 2D and E), similar to FEF_SEM_ neurons. Further, the preparatory-related modulation of activity in NRTP and DLPN encodes expectation of target speed in a fashion like FEF_SEM_: larger modulation of activity during the fast-context compared to the slow-context (Figures 2G, H, J, and K). Statistics confirmed the impression from the graphs. In NRTP, preparatory modulation of activity in the fast-versus slow-context was 17.68 +/- 0.12 spikes/s versus 13.10 +/- 0.08 spikes/s (mean +/- s.e.m.; paired sample t-test, p = 2.14 × 10^−10^, n = 182, Cohen’s d = 0.50). In DLPN, preparatory modulation of activity in the fast-versus slow-context was 13.57 +/- 0.24 spikes/s versus 9.89 +/- 0.16 spikes/s (mean +/- s.e.m.; paired sample t-test, p = 0.01, n = 66, Cohen’s d = 0.32).

### Motor preparation modulates visual motion processing in pursuit-related pontine areas

Motor preparation modulates the gain of visual-motor transmission, leading to the question of whether it also influences visual motion processing in the pons. To evaluate the state of gain of the pursuit system without adjusting gain and understand how visual-motor gain related preparatory signals influence visual motion processing in NRTP and DLPN, we probed visual-motor gain with brief, 50-ms pulses of 5 deg/s visual motion during fixations early versus late in preparation (Figure 3A). Because the motion pulses were identical, differences in behavioral and neural responses to pulses at different times could only be due to changes in visual-motor gain and its influence on visual motion signals.

**Figure 3.**
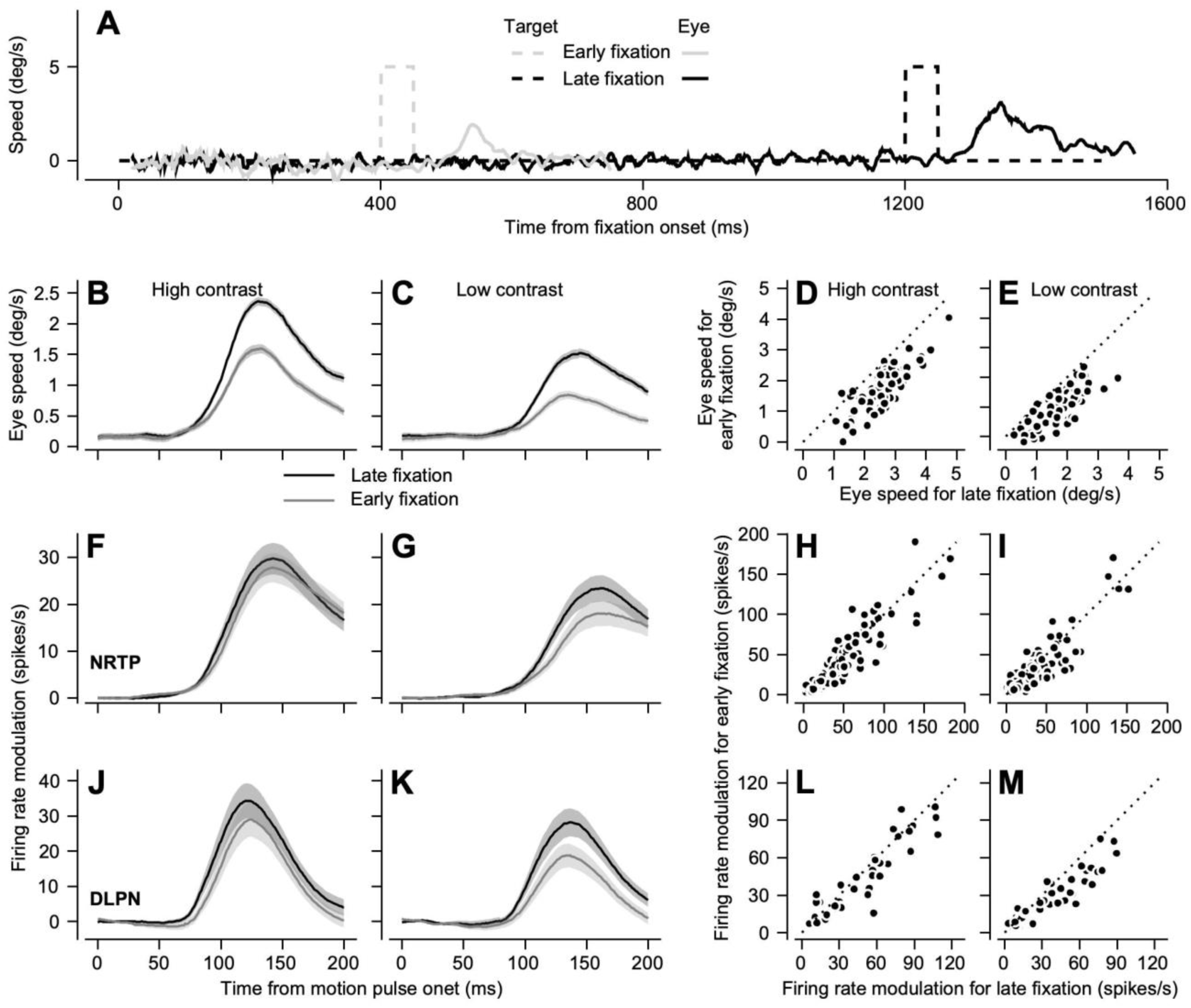
Fixation duration affects the gain of eye velocity and the firing of NRTP and DLPN neurons in response to pulses of visual motion. **A**: Timing of stimuli within a trial. Dashed and continuous traces show target and eye velocity; gray and black eye velocity traces show responses for pulses of visual motion early versus late in fixation. **B**: Black and grey curves plot the eye speed responses over time averaged across experiments for pulses of visual motion for high-contrast targets delivered during late versus early in fixation. **C**: Same as **B**, for low-contrast targets. **D:** Symbols plot the peak value of the eye speed response in individual experiments to pulses of visual motion of high-contrast targets delivered late versus early in fixation. **E**: Same as **D** for low-contrast targets. **F:** Black and grey curves plot the population-averaged firing rate modulation responses of NRTP neurons to pulses of visual motion of high-contrast targets delivered late versus early in fixation. **G**: Same as **F** for low-contrast targets. **H**: Symbols plot the firing rate modulation of individual neurons in NRTP to pulses of visual motion of high-contrast targets delivered late versus early in fixation. **I**: Same as **H** for low-contrast targets. **J**: Same as **F** for DLPN. **K**: Same as **G** for DLPN. **L**: Same as **H** for DLPN. **M**: Same as **I** for DLPN. Dotted lines plot the unity line. Error-bands represent S.E.M.

Visual-motor gain increases during fixation as time draws nearer to the expected onset of target motion. Consistent with previous results, eye speed responses were larger when pulses of motion were delivered late during fixation compared to when they were delivered early; responses also were larger for motion of high-contrast versus low-contrast targets (Figures 3B and C). The consistency of the effects on eye movement responses across many recordings can be appreciated from the scatter plots in Figures 3D and E. Therefore, we can use pulses of visual motion at different times during fixation to study how the state of visual-motor gain influences visual motion processing in NRTP and DLPN.

Motor preparation enhances responses to visual motion in both NRTP and DLPN. Pulses of visual motion caused larger modulation of firing rate when delivered later compared to earlier during fixation (Figures 3F and G). These trends were consistent across individual neurons, as shown in scatter plots of the response to a pulse of motion early versus late in fixation (Figures 3H, I, L, M). In both structures and for the motion of both high-contrast and low-contrast targets, the effect of fixation time was statistically significant, usually at very high levels (Table 1A). In general, the modulations of response amplitude by fixation time were somewhat larger for the motion of low-versus high-contrast targets. But the absolute eye speed responses and neural responses for pulses of motion delivered during early or late fixation were larger for high-contrast compared to low-contrast motion (Table 1C).

**Table 1:** Summary of statistics for responses to brief pulses of target motion at different times, using different target contrasts, and in different speed contexts. Entries show mean values across the sample population with standard error of the mean, p-value from paired t-tests on the full sample and the number of data points, and Cohen’s d to quantify effect size.

| **A. Effect of fixation time, speed contexts combined** |  |  |  |  |  |  |  |  |  |
| --- | --- | --- | --- | --- | --- | --- | --- | --- | --- |
|  | High contrast, late versus early pulses |  |  |  |  | Low contrast, late versus early pulses |  |  |  |
|  | Late | Early | p-value, n | Cohen's d |  | Late | Early | p-value, n | Cohen's d |
| Eye speed (deg/s) | 2.63<br>+/-0.07 | 1.77<br>+/-0.07 | $5.94 \times 10^{-32}$<br>n = 86 | 2.02 | | 1.78<br>+/-0.07 | 1.02<br>+/-0.06 | $1.82 \times 10^{-30}$<br>n = 86 | 1.92 |
| DLPN (spikes/s) | 47.07<br>+/-4.87 | 41.40<br>+/-4.59 | 0.0054,<br>n = 39 | 0.47 | | 38.94<br>+/-4.14 | 29.15<br>+/-3.07 | $3.31 \times 10^{-6}$<br>n = 39 | 0.87 |
| NRTP (spikes/s) | 43.89<br>+/-3.21 | 39.45<br>+/-3.18 | 0.0013,<br>n = 123 | 0.30 | | 35.60<br>+/-2.67 | 29.42<br>+/-2.60 | $1.01 \times 10^{-5}$<br>n = 123 | 0.42 |
| **B. Effect of speed context, late pulses** |  |  |  |  |  |  |  |  |  |
|  | High contrast, fast versus slow context |  |  |  |  | Low contrast, fast versus slow context |  |  |  |
|  | Fast | Slow | p-value, n | Cohen's d |  | Fast | Slow | p-value, n | Cohen's d |
| Eye speed (deg/s) | 2.96<br>+/-0.08 | 2.35<br>+/-0.07 | $3.3 \times 10^{-23}$<br>n = 83 | 1.52 | | 2.08<br>+/-0.08 | 1.59<br>+/-0.07 | $4.5 \times 10^{-18}$<br>n = 83 | 1.22 |
| DLPN modulation (spikes/s) | 49.50<br>+/-4.82 | 50.17<br>+/-4.95 | 0.64<br>n = 38 | N/A |  | 42.75<br>+/-4.43 | 41.59<br>+/-4.16 | 0.59<br>n = 38 | N/A |
| NRTP modulation (spikes/s) | 46.76<br>+/-3.27 | 45.84<br>+/-3.28 | 0.36<br>n = 122 | N/A |  | 38.60<br>+/-2.70 | 37.46<br>+/-2.75 | 0.24<br>n = 122 | N/A |
| NRTP absolute (spikes/s) | 86.50<br>+/-5.44 | 82.38<br>+/-5.29 | $2.63 \times 10^{-5}$<br>n = 122 | 0.40 | | 77.92<br>+/-5.05 | 73.62<br>+/-4.75 | $7.12 \times 10^{-5}$<br>n = 122 | 0.37 |
| **C. Effect of contrast** |  |  |  |  |  |  |  |  |  |
|  | Late pulse, contexts combined |  |  |  |  | Early pulse, contexts combined |  |  |  |
|  | High | Low | p-value, n | Cohen's d |  | High | Low | p-value, n | Cohen's d |
| Eye speed (deg/s) | 2.63<br>+/-0.07 | 1.78<br>+/-0.07 | $1.6 \times 10^{-34}$<br>n = 86 | 2.20 | | 1.77<br>+/-0.07 | 1.02<br>+/-0.06 | $5.1 \times 10^{-30}$<br>n = 86 | 1.89 |
| DLPN (spikes/s) | 47.07<br>+/-4.87 | 38.94<br>+/-4.14 | $6.3 \times 10^{-5}$<br>n = 39 | 0.72 | | 41.40<br>+/-4.59 | 29.15<br>+/-3.07 | $3.08 \times 10^{-6}$<br>n = 39 | 0.88 |
| NRTP (spikes/s) | 43.89<br>+/-3.21 | 35.60<br>+/-2.67 | $2.9 \times 10^{-12}$<br>n = 123 | 0.70 | | 39.45<br>+/-3.18 | 29.42<br>+/-2.60 | $2.3 \times 10^{-11}$<br>n = 123 | 0.66 |
|  | Fast context, late pulses |  |  |  |  | Slow context, late pulses |  |  |  |
|  | High | Low | p-value, n | Cohen's d |  | High | Low | p-value, n | Cohen's d |
| Eye speed (deg/s) | 2.96<br>+/-0.08 | 2.08<br>+/-0.08 | $3.5 \times 10^{-31}$<br>n = 83 | 2.04 | | 2.35<br>+/-0.07 | 1.59<br>+/-0.07 | $2.2 \times 10^{-27}$<br>n = 83 | 1.79 |
| DLPN (spikes/s) | 49.50<br>+/-4.82 | 42.75<br>+/-4.43 | $8.9 \times 10^{-4}$<br>n = 38 | 0.59 | | 50.17<br>+/-4.95 | 41.59<br>+/-4.16 | $4.0 \times 10^{-4}$<br>n = 38 | 0.63 |
| NRTP (spikes/s) | 86.50<br>+/-5.44 | 38.60<br>+/-2.70 | $6.61 \times 10^{-9}$<br>n = 122 | 0.57 | | 45.84<br>+/-3.28 | 37.46<br>+/-2.75 | $5.0 \times 10^{-10}$<br>n = 122 | 0.61 |

Our data contain an unplanned behavioral control that provides a good test case for how well the neural responses in DLPN and NRTP encode movement. Eye speed response magnitudes are essentially the same for high-contrast pulses of visual motion delivered during early fixation and low-contrast pulses of visual motion delivered late during fixation (paired sample t-test, p = 0.82, n = 86). Consistent with the behavioral data, there was no difference in neural responses in DLPN for the same comparison: high-contrast visual motion pulses delivered early during fixation versus low-contrast visual motion pulses delivered late during fixation (paired sample t-test, p = 0.25, n = 39). In NRTP, however, neural responses to high-contrast visual motion pulses delivered early during fixation were larger than those to low-contrast visual motion pulses delivered during late fixation (paired sample t-test, p = 0.0052, n = 123, Cohen’s d = 0.26). Thus, responses in DLPN closely resemble a motor command while responses in NRTP also depend on some movement-independent features of contrast.

### Speed context does not influence visual motion processing in pursuit-related pontine areas

Expectation of the upcoming target speed controls the amplitude of preparatory activity in FEF_SEM,_ NRTP, and DLPN (Figure 2F-H). Higher preparatory activity leads to higher gain of visual-motor transmission during fixations that lead up to expected target motion. In agreement with previous data, eye speed responses to pulses of visual motion were larger during the fast compared to the slow context in the experiments reported here (Figures 4A-D). Mean eye speeds across recordings, standard errors of the mean, and other statistics appear in Table 1B.

**Figure 4.**
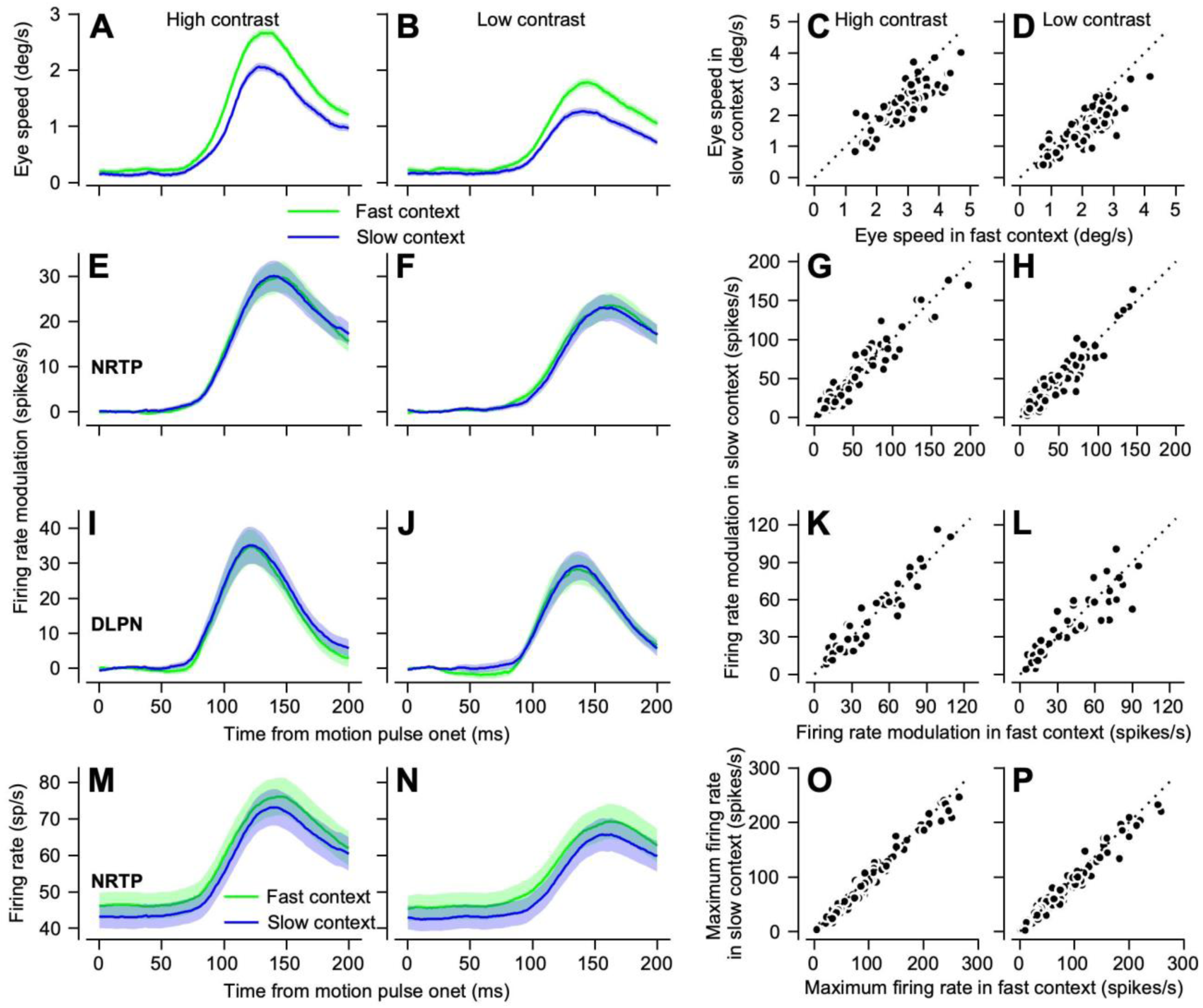
Speed context dissociates responses of NRTP and DLPN neurons from eye velocity responses to pulses of visual motion. **A**: Red and black curves plot eye speed responses over time averaged across experiments to pulses of visual motion of high-contrast targets delivered during the fast versus slow contexts. **B**: Same as **A** for low-contrast targets. **C**: Symbols plot the maximum value of the eye speed response for individual experiments to pulses of visual motion of high-contrast targets delivered during the fast versus slow speed contexts. **D**: Same as **C** for low-contrast targets. **E:** Red and black curves plot the population-averaged firing rate modulation over time of NRTP neurons to pulses of visual motion of high-contrast targets delivered during the fast versus slow contexts. **F**: Same as **E** for low-contrast targets. **G**: Symbols plot the firing rate modulation of individual neurons in NRTP to pulses of visual motion of high-contrast targets delivered during the fast versus slow contexts. **H**: Same as **G** for low-contrast targets. **I**: Same as **E** for DLPN. **J**: Same as **F** for DLPN. **K**: Same as **G** for DLPN. **L**: Same as **H** for DLPN. **M:** Same as **E** but plotting absolute firing rate for high-contrast targets in NRTP. **N**: Same as **M** for low-contrast targets. **O:** Same as **G** but plotting absolute firing rate for high-contrast targets in NRTP. **P**: Same as **O** for low-contrast targets. Dotted lines plot the unity line. Error-bands represent S.E.M.

The change in preparatory activity based on expectation of target speed (Figure 4) did not influence visual-motor processing in NRTP and DLPN in the same way that fixation time did. The modulation of preparatory activity by speed expectation had no effect on visual motion related responses in neither NRTP nor DLPN. There were no statistical differences across the population responses of both areas to visual motion pulses delivered during the fast versus slow context, for high- or low-contrast targets (Figures 4E-L). Mean responses across neurons, standard errors of the mean, and other statistics appear in Table 1B. Again, behavioral and neural responses were larger in both NRTP and DLPN for the motion of high-compared to low-contrast targets within both speed contexts (Table 1C).

Because the statistically indistinguishable visual motion related responses are riding on top of different settings of preparatory activity in the different speed contexts, there was a difference in the absolute firing of NRTP neurons during the response to visual motion pulses. Consistent with larger modulation of preparatory activity during the fast context, the firing rate of NRTP neurons was slightly higher throughout the entire response to the pulses of both high- and low-contrast visual motion (Figures 4M-P). Mean responses across neurons, standard errors of the mean, and other statistics appear in Table 1B.

A similar analysis shows that speed context does affect the responses of neurons in NRTP and DLPN during pursuit initiation (Supplementary Figure 1).

### Neural activity in NRTP and DLPN can be linked directly to the behavioral effects of fixation time and contrast

The preceding sections established that the differences in neural responses to visual motion pulses across motor preparation time and visual motion contrast are correlated with the gain of visual-motor transmission as probed in eye movement behavior. We now shift our efforts to establishing a more direct link between the neural and behavioral responses to visual motion pulses. First, we fit a linear model between the neural responses of each member of our population of neurons in NRTP and DLPN and the behavioral results. Second, we compare the relative effect sizes of the neural and behavioral responses.

The sum of a linear weighting of neural responses across our population of NRTP and DLPN neurons fits nicely with the effects of fixation time and contrast on behavioral responses to visual motion pulses. To generate the linear weighting, we normalized each neuron to its range of firing rate across both fixation times and both contrasts and then used regularized regression to compute a weight vector linking each neuron’s response to evoked eye speed. We created separate linear models for DLPN and NRTP and both predicted the eye movements almost perfectly (Figures 5A and D). Importantly, the linear model weights were distributed smoothly across each population of neurons (Figures 5C and F), meaning that the regression did not simply select a small handful of neurons that were best related to the behavioral effects.

**Figure 5.**
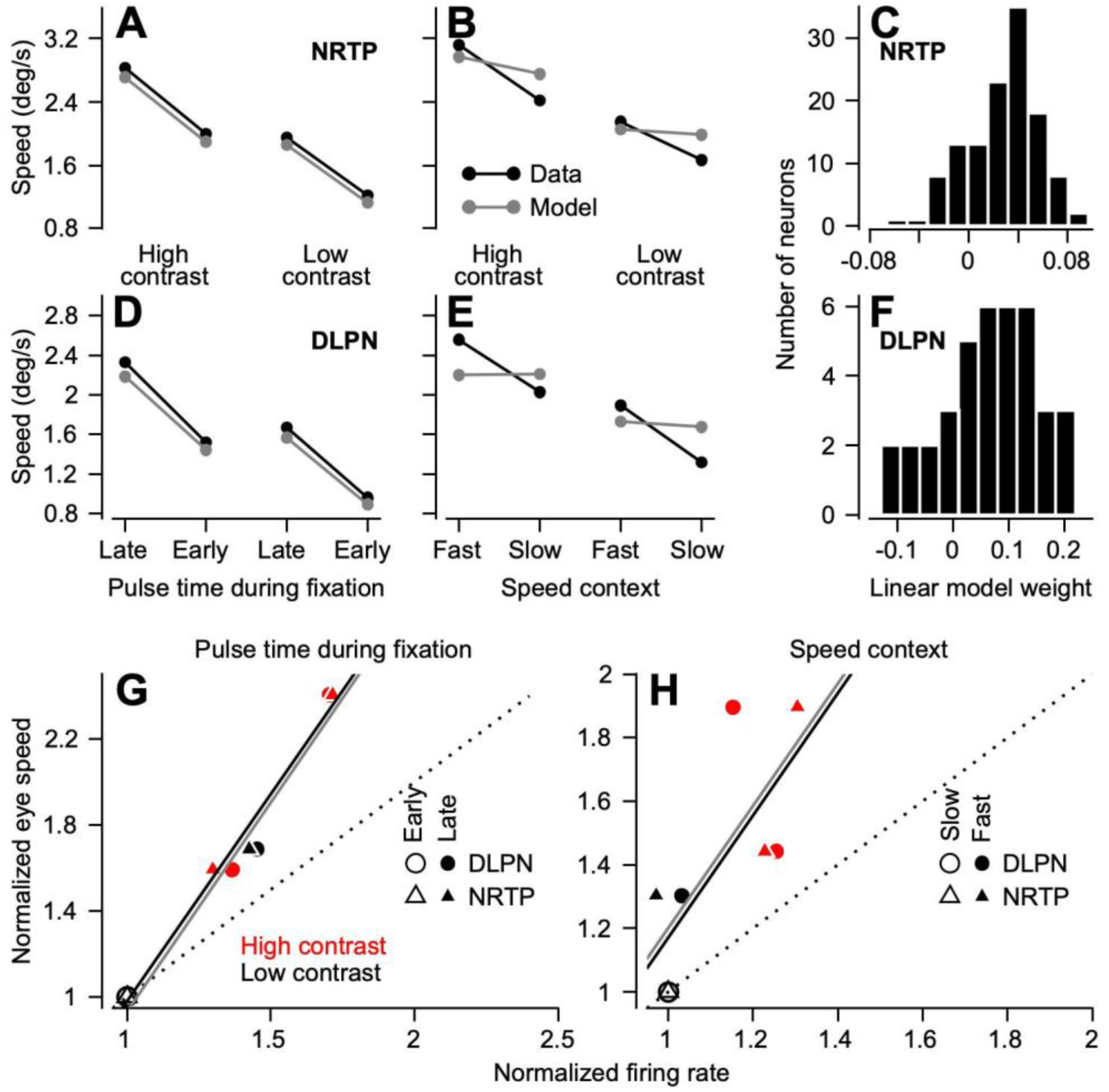
A linear model demonstrates quantitative links between the neural and behavioral effects of fixation duration and contrast. **A**: Black and grey symbols connected by lines compare the average behavioral responses for fixation times and target contrasts to the predictions of a model based on a linear weighting of responses in NRTP. **B:** Same as **A** but comparing average behavioral responses to the predictions of the linear model for different speed contexts and target contrasts. **C**: The distribution of weights used for different NRTP neurons in the linear model. **D**: Same as **A** for DLPN. **E**: Same as **B** for DLPN. **F**: Same as **C** for DLPN. **G:** Filled and open symbols plot normalized average eye speeds as a function of normalized average firing rates of DLPN (circles) and NRTP (triangles) neurons in response to pulses of visual motion of high-contrast (red) and low-contrast (black) targets delivered late versus early in fixation. Black and grey solid lines plot the best fit line for the NRTP and DLPN data. Dotted lines plot the unity line. **H**: Same as **G** but comparing data for the fast and slow contexts.

The linear model provides further evidence that the behavioral effects of speed context are not represented in the modulation of activity of NRTP and DLPN. The models that we generated to predict the behavioral effects of fixation duration and visual motion contrast failed to predict the behavioral effects of speed context (Figures 5B and E; compare Fast versus Slow). The models did, however, predict the difference between behavioral responses to high- and low-contrast visual motion pulses in the speed context data (Figures 5B and E; compare left and right pairs of symbols).

Finally, we demonstrate that the neural effects, while able to predict the behavioral responses through a linear model, have a smaller magnitude than the behavioral effects meaning that some form of amplification occurs downstream. For the experiments that varied the time of the pulses, we normalized the neural and behavioral data relative to the values for the responses to motion pulses of low contrast stimuli delivered early in fixation. For the experiments that varied speed context, we normalized relative to the values for visual motion of low-contrast targets in the slow-speed context. For each set of experiments, we plotted the normalized eye speed as a function of normalized neural responses and fitted a line to the data.

We find a strong linear relationship between the neural and behavioral effects of fixation duration and visual motion contrast in both NRTP and DLPN (Figure 5G), but the slope of the line is not one, meaning that the relative neural effects underestimate the relative magnitude of the behavioral effects. The significance of these slopes and their high r^2^ values suggest that the firing rates are very linear across the behavioral effects of fixation duration and contrast. For NRTP, the slope for the line is 1.93 (Figure 5G, black line; p = 0.0083, r^2^ = 0.98). For DLPN, the slope for the line is 1.94 (Figure 5G, gray line; p = 0.017, r^2^ = 0.97). However, the fact that the slope is greater than one suggests that there is further computation downstream and/or that the integration of descending motor commands is not perfectly linear.

In contrast, the linear fit between the neural and behavioral effects of speed context is poor in both NRTP and DLPN. For NRTP, the slope of the fit is 1.92, but is not significantly different from zero and accounts for a much smaller amount of the variance (Figure 5H, black line; p = 0.14, r^2^ = 0.73). For DLPN, the slope of the fit is 1.92 but is not statistically significant and accounts for even less of the variance (Figure 5H, gray line; p = 0.40, r^2^ = 0.36). Qualitative inspection of Figure 5H reveals that the variance captured by these regression lines is mostly the variance across contrast and not across speed context, consistent with the idea that the behavioral effects of target contrast are represented in NRTP and DLPN.

### Visual motion signals in the pons encode stimulus speed and contrast as a rate code

A long-standing question for sensory-motor coding across systems in general and for pursuit specifically is how the place code for visual motion speed in area MT is converted to a rate code to drive movement (Groh, 2001). Transformation to a rate code is necessary to account for the fact that faster targets need to elicit larger changes in the activity of motoneurons to drive faster eye movements. The doctrine in the field has been that the transformation is based on finding the preferred speed at the peak of the MT population response, that vector averaging is an appropriate decoding computation, and that the result should be contrast invariant (Lisberger, 2010). Our data challenge all these assumptions.

Neurons in both NRTP and DLPN responded strongly (Figure 6) to large field visual motion with a small central scotoma during fixation (see Methods for details). The population PSTHs show responses that scale monotonically with visual motion speed (Figures 6A, B, C, and D; see Supplementary Figure 2 for a comparison of responses to passive visual motion versus during active pursuit initiation). The normalized speed tuning curves for each individual neuron in our populations of neurons in DLPN (Figures 6E and F) and NRTP (Figures 6G and H) show that the majority encode speed monotonically. Firing rate increases as a function of speeds. In the place code in MT, in contrast, almost all neurons show responses that are tuned for speed with peak firing across the range of stimulus speeds we presented.

**Figure 6.**
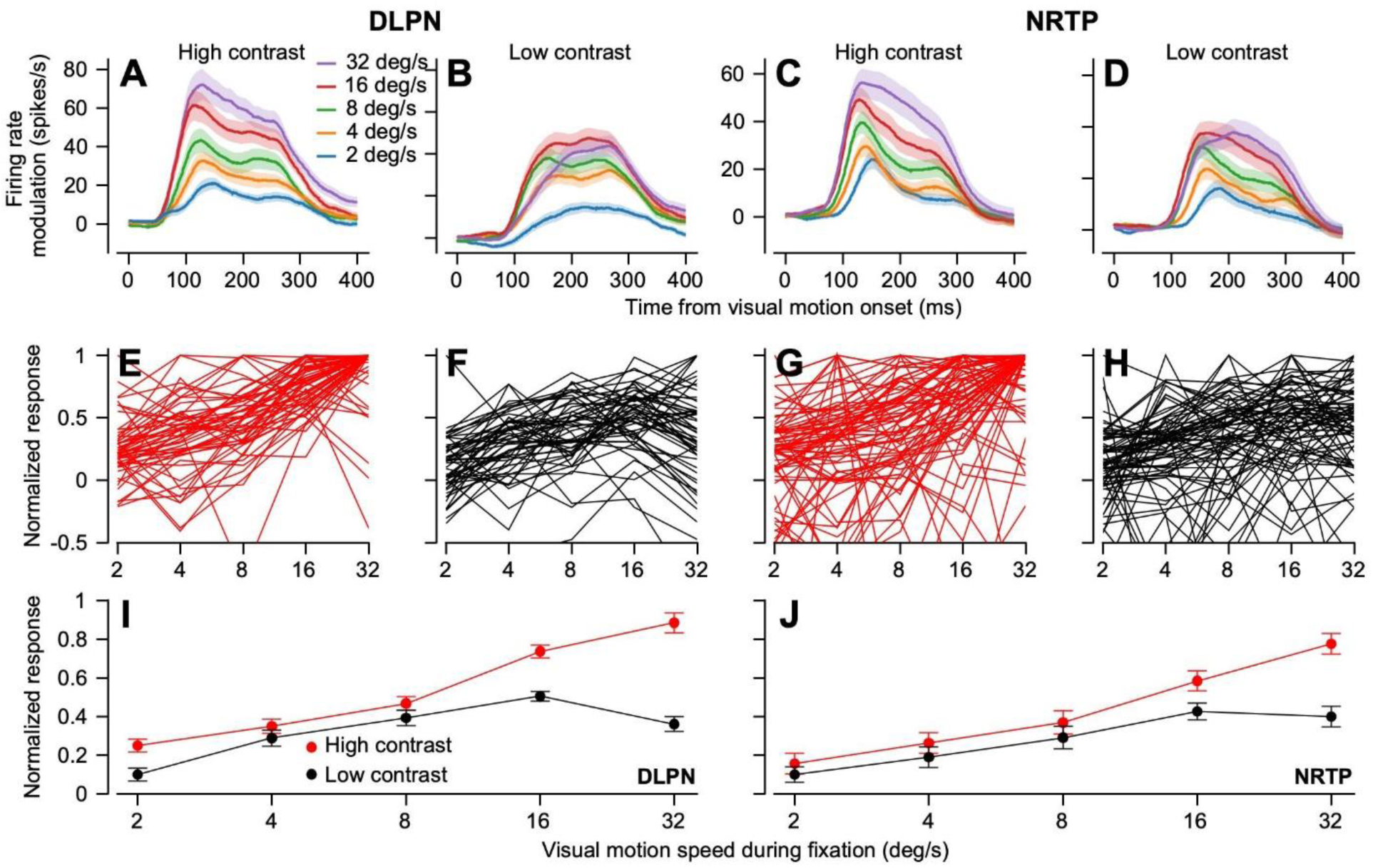
DLPN and NRTP neurons respond vigorously to passive visual motion that is not associated with eye motion. **A:** Different colored traces plot the population-averaged firing rate modulation of DLPN to passive visual motion of high-contrast targets delivered at different speeds. **B**: Same as **A** for low-contrast targets. **C**: Same as **A** for NRTP. **D**: Same as **B** for NRTP. **E**: Traces show firing rate modulation of DLPN neurons normalized to the maximum firing rate across conditions plotted as a function of visual motion speed for high-contrast targets. Each curve shows responses for a single neuron. **F**: Same as **E** for low-contrast targets. **G**: Same as **E** for NRTP. **H**: Same as **F** for NRTP. **I**: Red and black symbols connected by lines show the normalized population average speed tuning of DLPN neurons for high-contrast versus low-contrast targets. **J**: Same as **I** for NRTP.

The representation of stimulus speed is not contrast-invariant in the pons and the contrast-variance agrees with that for the initiation of pursuit. Averaged across our population of DLPN (Figure 6I) and NRTP (Figure 6J) neurons, responses are generally larger for the motion of high-versus low-contrast stimuli (red versus black symbols). The impact of contrast is greater for faster visual motion speeds, consistent with the behavioral findings and responses during the initiation of pursuit in Figure 1. Thus, at least some of the behavioral effects of stimulus contrast can be explained by the way that the pons decodes speed from the visual motion signals from area MT.

Figure 6 demonstrates how target speed is *encoded* in DLPN and NRTP. The important related question is how these two sites in the pursuit circuit *decode* visual motion speed from the population response of area MT. Previous models have assumed some form of vector averaging or divisive normalization that would predict contrast-invariant responses to the same speed (Churchland & Lisberger, 2001; Priebe et al., 2001; Priebe & Lisberger, 2004). Our data show that the responses are not contrast-invariant and we find that a much simpler decoder may be adequate.

We start with a model of area MT responses based on well-documented effects of speed and contrast on the neural responses of MT neurons (Krekelberg et al., 2006; Nover et al., 2005). Each of 60 model MT neurons shows responses that are tuned for stimulus speed (Figure 7A) with a log-uniform distribution of preferred speeds (Figure 7B). Reductions in stimulus contrast reduce the amplitude of the tuning curves and shift their peaks to the left because contrast has a larger effect on the response to faster versus slower speeds (Figure 7A). We create a population response for each target speed and contrast by reading out each neuron’s response for the selected conditions. We then fit a linear model for each of NRTP and DLPN that transforms the 10 population responses (5 speeds x 2 contrasts) for area MT into the measured responses of DLPN and NRTP across visual motion speeds and contrast. The simple linear decoding model, with different weights for each structure, captures the visual motion responses that we recorded in NRTP (Figures 7D and E) and DLPN (Figures 7G and H). The weight vectors (Figures 7C and F) reveal that the linear model weighted model units with faster preferred speeds more strongly. Thus, the decoder of responses from area MT to drive pursuit may be much simpler to implement with neurons than previously believed.

**Figure 7.**
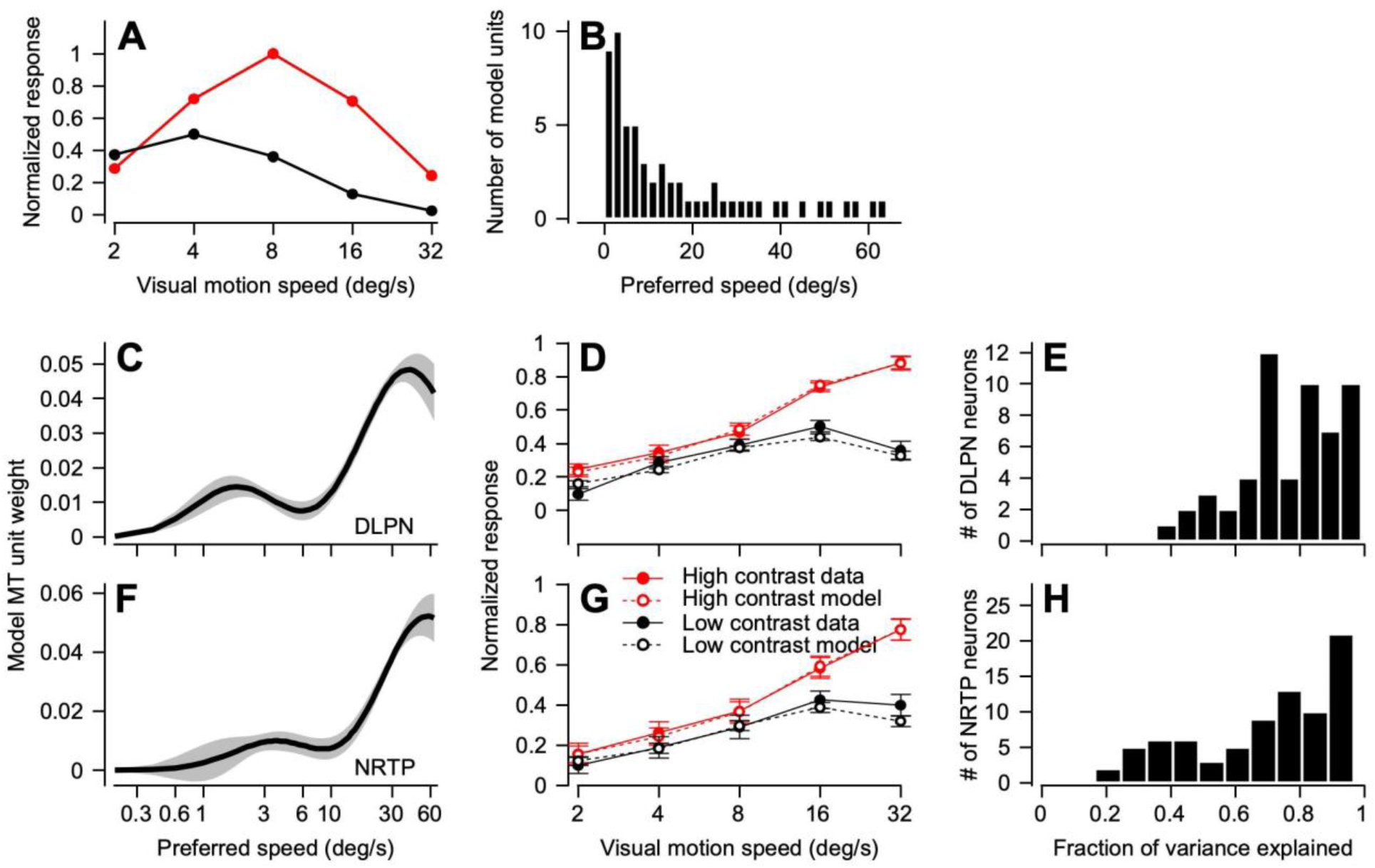
A linear decoder transforms the place code in area MT to a rate code in NRTP and DLPN across target speeds and contrasts. **A**: Red and black symbols connected by lines plot the simulated speed-tuning of an example model MT unit with preferred speed of 8 deg/s for passive motion of high-contrast versus low-contrast targets. **B**: Distribution of preferred speeds across all the model units in our model population of area MT. **C**: Average weights used in the linear model as a function of preferred speed to reproduce the speed tuning of the population of DLPN neurons across targets speeds from 2 to 32 deg/s and for high-contrast and low-contrast targets. Error-bands represent S.E.M. **D**: Filled and open symbols connected by continuous and dashed lines compare speed tuning for the data and the predictions of the linear model averaged across DLPN neurons. Red and black symbols and lines show data and predictions for high-contrast versus low-contrast targets. Error-bars represent S.E.M. **E**: distributions of variance explained by the linear model across our population of DLPN neurons. **F**: Same as **C** for NRTP. **G**: Same as **D** for NRTP. **H**: Same as **E** for NRTP.

## Discussion

In the present paper we take a major step towards understanding how the outputs from multiple areas of the cerebral cortex are combined in the sub-cortical motor system to form a motor command. We can take this essential step in the smooth pursuit eye movement system because of three advantages of the system. (i) We know the cortical areas that are involved and their primary anatomical projections to the brainstem. (ii) We already have partitioned the conceptual smooth pursuit system into multiple parallel components and assigned those components to specific cortical areas. (iii) The brainstem and cerebellar components of the smooth pursuit system are known and enough information is available to allow us to interpret the discharge of neurons in terms of their causal effect on motoneuron firing and eye movement. Thus, the subcortical processing of smooth eye movements provides an exemplar that generalizes to the bigger question of how cortical representations of movement are transformed into commands for muscle activity and movement.

In the smooth pursuit system, we start from two fundamental concepts that drive our analysis of sub-cortical mechanisms. First, visual-motion drives pursuit, but it does not have unchecked access to the final motor pathways. It is subject to modulation of the strength, or “gain”, of visual-motor transmission (Schwartz & Lisberger, 1994). Gain is controlled by the output of one key cortical region, FEF_SEM_ (Tanaka & Lisberger, 2001). Implementation of gain control on visual motion signals probably occurs sub-cortically at the convergence of outputs from area MT and FEF_SEM_. The data in the present paper provide a major advance by showing that the site and mechanism of gain control lies partly in or before NRTP and DLPN, and partly downstream. Second, the speed of visual motion is represented in MT, but as a “place code” that must be converted to a “rate code” to drive pursuit (Priebe & Lisberger, 2004). Our data reveal that the transformation is complete in the firing of pontine neurons, and that it can occur through a weighted sum of the firing of MT neurons. The requisite decoder could be constructed neurally simply by proper setting of synaptic strengths in the outputs from MT to the pons.

### Motor preparation in the pursuit system affects visual motion processing in the pons

We use motor preparation to study the neural mechanisms of gain control. In the pursuit system, motor preparation modulates the state of visual-motor gain (Kodaka & Kawano, 2003; Tabata et al., 2006). Preparatory enhancement of visual-motor gain has been linked to FEF_SEM_ neurons, which display preparatory ramps of firing during fixation that leads up to pursuit (Darlington & Lisberger, 2020; Mahaffy & Krauzlis, 2011; Tanaka & Fukushima, 1998). Expectation of faster target motion causes larger amplitudes of preparatory activity and larger behavioral readouts of visual-motor gain (Darlington et al., 2018; Darlington et al., 2017).

The presence of preparatory activity in pontine neurons demonstrates that preparatory signals are part of the output of FEF_SEM_ and therefore could modulate pontine responses to visual motion inputs. Indeed, the responses to pulses of target motions at different times indicate that the preparatory signals modulate the excitability of NRTP and DLPN for visual motion inputs from MT/MST. Pontine visual responses are enhanced as motor preparation progresses; they do not simply ride on top of preparatory-related changes in baseline firing.

### What is computed locally in the pons?

It is important to distinguish between features of pontine responses that can be attributed to upstream areas like FEF_SEM_ and area MT, and features that are a direct result of local computations in the pons. Anatomically, FEF_SEM_ is the primary input to NRTP and area MT/MST is the primary input to DLPN (Brodal, 1980a; Distler et al., 2002; Giolli et al., 2001; May & Andersen, 1986). Consistent with the anatomical findings, previous neurophysiological experiments using only a basic pursuit task showed agreement between the signals in FEF_SEM_ and NRTP and the signals in area MT/MST and DLPN (Mustari et al., 2009; Ono et al., 2005).

The present paper provides strong evidence that these cortico-pontine pathways aren’t just relays – they transform their inputs. Consider DLPN. First, the presence of preparatory activity in DLPN implies that some of the output from FEF_SEM_ propagates either directly or indirectly to DLPN. Preparatory activity in DLPN neurons encodes speed context similarly to FEF_SEM_ neurons: larger modulation of activity during the fast-compared to the slow-speed context. To our knowledge, preparatory modulation of activity has not been demonstrated in either area MT or MST. Third, modulation of visual-motor gain is not represented in area MT. Responses to pulses of visual motion delivered early versus late during fixation are identical (J.P. Mayo and S.G. Lisberger, unpublished observations). Taken together, these findings suggest that visual-motor gain signals from FEF_SEM_ modulate the processing of parallel visual motion inputs locally within DLPN.

What is computed locally within NRTP is less clear. Most of the features of our NRTP data are consistent with what has been previously shown in FEF_SEM_. The preparatory activity in NRTP agrees with that found in FEF_SEM_. Furthermore, visual motion related signals have been reported previously in FEF_SEM_ (Bakst et al., 2017; Ono & Mustari, 2009). Indeed, FEF_SEM_ activity during pursuit initiation can be modeled as a Bayesian-like estimation of target speed that combines preparatory signals with visual motion input (Darlington et al., 2018). Still, we cannot draw a definitive conclusion about local versus inherited modulation of visual responses in NRTP because no experiment has delivered pulses of visual motion during motor preparation while recording in FEF_SEM_.

### Further integration of visual motion and visual-motor gain occurs downstream of the pons

Our data reveal that certain transformations need to happen downstream from DLPN and NRTP. For example, both NRTP and DLPN neurons show large responses to passive visual motion presented during fixation when there is no smooth eye movement. Both also show ramps of preparatory activity during fixation without smooth eye movement. Perhaps the large pursuit-related output from FEF_SEM_ modulates the output from DLPN and NRTP in downstream circuits. As an alternative, fixation signals in the brainstem might prevent eye movement by gating the large field visual motion and preparatory responses. For example, omnipause neurons in the pontine and medullary reticular formation are part of the circuit that generates bursts of activity to drive saccades (Cohen & Henn, 1972; Evinger et al., 1982; Keller, 1974). Just as they inhibit movement-driving bursts of firing until it is time to initiate the saccade, omnipause neurons might be well-suited to suppress smooth eye movements to visual motion during fixation. They have suggestive response features during smooth pursuit but need to be studied in relation to visual-motor gain and visual motion signals (Missal & Keller, 2002).

The effects of the time of the motion pulse during fixation and the contrast of the target on the firing rate responses in DLPN and NRTP underestimate the relative magnitude of the behavioral effects by approximately a factor of 2. Therefore, further integration of motor preparation signals and visual motion may occur downstream of DLPN and NRTP. As an alternative, downstream neural mechanisms may create a non-linear transformation in the mapping from pontine responses to motor commands and motor output. For example, moving the eye in the pulling direction of a muscle recruits silent extraocular motoneurons as the eye position crosses their thresholds (Fuchs et al., 1988; Robinson, 1970). Recruitment could amplify a relatively smaller change in the motor command and cause a larger-than-expected eye movement.

Finally, the responses of pontine neurons imply that the integration of speed context related visual-motor gain information with visual motion occurs downstream of NRTP and DLPN. Expectation of fast versus slow target motion does not modulate pontine responses to brief pulses of visual motion but does affect the size of eye speed responses. We note, however, that the firing rate response to visual motion in NRTP rides on top of different baseline firing rates set by the preparatory response in the fast and slow context. Therefore, the firing rate coming out of NRTP is higher throughout the response to the pulse of visual motion during the fast context. This could, for example, lead to recruitment of a larger pool of motor neurons.

### Transforming a sensory place code into commands for movement

We find that the computation for reading out a command for pursuit eye movement from MT can be very simple. A linear, weighted sum of MT responses reproduces the response of NRTP and DLPN neurons as a function of both target speed and contrast. We performed this analysis for passive visual motion during fixation to avoid the complication of gain-control modulation during pursuit initiation, but we obtain very similar effects of both speed and contrast during pursuit initiation. Therefore, we think that the sensory place code can be transformed into motor commands by differential weighting of neurons with different preferred speed. It does not require more complex computations such as vector averaging through response normalization (Priebe & Lisberger, 2004).

### Concluding thoughts about sub-cortical mechanisms of motor control

Our data suggest a revision in thinking about how sub-cortical motor circuits process cortical signals to generate commands for movement. First, we must abandon the traditional thought that the pontine nuclei are “relay nuclei” in the cortico-ponto-cerebellar pathways. Considerable processing occurs in the pontine nuclei. Second, we need to alter thoughts about localization of functions such as control of the gain of visual-motor transmission. The interaction of parallel descending cortical signals seems to be distributed across sub-cortical areas. Distributed integration of visual motion signals and visual-motor gain probably should have been expected given that the outputs of NRTP and DLPN are distributed relatively privately to the oculomotor vermis versus the flocculus and paraflocculus (Brodal, 1980b, 1982; Glickstein et al., 1994). Third, we need to simplify how we think about using the place code in sensory areas to drive motor output. The effects of both target speed and contrast on the visual responses on pontine neurons can be attributed to a simple weighted sum of the firing of MT neurons. Finally, we may need to adjust conclusions about separation of the final motor pathways for different kinds of movements. For example, the inhibitory omnipause neurons in the saccadic motor pathways may be ideally suited to also prevent smooth eye movements during fixation by gating preparatory activity and passive visual responses that emerge from DLPN and NRTP.

## Acknowledgements

We thank Stefanie Tokiyama, Bonnie Bowell, Scott Ruffner, and Steven Happel for technical assistance. We thank David Herzfeld for providing the code for the Full Binary Pursuit spike sorter in Julia. Research supported by R01-EY027373 (SGL) and F30-EY027684 (TRD).

## Methods

### Experimental model and subject details

We conducted experiments on two male rhesus monkeys aged 11-14 years and weighing 10-13 kg. Each monkey was surgically implanted with a post on the skull to restrain head movement and a scleral coil to track eye movements (Fuchs & Robinson, 1966). After recovering from surgery, monkeys were trained to smoothly track moving visual targets displayed on a 24-inch CRT monitor with a refresh rate of 80 Hz that was positioned 35 cm from the monkey’s eyes. Horizontal and vertical components of eye position were sampled and stored at 1 kHz from the analog signals produced by the scleral search system. After the monkeys were fully trained, we performed a craniotomy and implanted a sealable recording cylinder over the craniotomy under stereotactic guidance: a chamber with 20-30 degrees of lateral tilt was aimed at 3.5 mm anterior, 1.0 mm lateral, and 8.0 mm dorsal relative to ear bar zero, the nominal coordinates of the oculomotor nucleus (OMN). All experimental procedures received prior approval from the Duke *Institutional Animal Care and Use Committee* and were in compliance with the National Institutes of Health’s *Guide for the Care and Use of Laboratory Animals*.

### Neurophysiology

For daily recordings, we lowered a guide tube into the brain approximately 15-20 mm below the dura mater and then used a microdrive to extend a single electrode an additional 20-30 mm beyond the end of the guide tube. We located nucleus reticularis tegmenti pontis (NRTP) and the dorsolateral pontine nucleus (DLPN) in several steps. First, we found OMN based on the characteristic burst-tonic firing in relation to saccadic eye movements. Second, we adjusted the trajectory of the penetration medially/laterally until we encountered neurons that exhibited the previously described basic responses for NRTP and DLPN during passive visual motion stimuli and/or smooth pursuit eye movements. The successful recording sites were appropriately located in anatomical positions relative to OMN. We found NRTP on penetrations were only slightly lateral to or sometimes through OMN; we found DLPN on penetrations that were considerably more lateral (for ipsilateral DLPN) or medial (for contralateral DLPN) to OMN. We found NRTP and DLPN at least 4-6 mm deeper than OMN. We also verified recording locations anatomically and confirmed that they were in NRTP and DLPN. At the conclusion of the recording experiments, both monkeys were deeply anesthetized and trans-cardially perfused with 100 mM phosphate-buffered saline followed by a 4% paraformaldehyde solution. Frozen sections of brain were cut, mounted on slides, and stained for Nissl substance to allow visualization of electrode tracks.

In Monkey Xt, some of the neurophysiological data were collected with Thomas Recording electrodes lowered using the Thomas Recording Mini Matrix System and Leopold Funnel Adapter. Voltage signals were amplified, filtered, and digitized at a sampling rate of 40 kHz using the Plexon MAP system. The timing and waveform of action potentials that reached a specified threshold were stored for offline analysis. Spikes were assigned to single units using the Plexon Offline Sorter. The rest of the neurophysiological data presented in this paper were collected using single FHC electrodes lowered hydraulically using a Narishige Microdrive (MO-97). Voltage signals were amplified and digitized at a sampling rate of 40 kHz using the OmniPlex system. For these data, wideband voltage signals were stored for offline analysis in which the data were digitally filtered, and spikes were assigned to single units via the semi-automated “Full Binary Pursuit” spike-sorting algorithm (Hall et al., 2021).

### Behavioral paradigms

During our neurophysiological data collection, monkeys were rewarded for successful completion of discrete trials. We delivered a droplet of juice at the end of the trial for maintaining a gaze position within a 4 × 4 degree window from the center of the target throughout the entire trial. Targets in all experiments were displayed against a grey background.

#### Visual motion pulse experiments

Trials were divided into blocks with strategic blends of target speeds. During a fast context block, the pursuit target moved at 20 deg/s for 80% of the trials and 10 deg/s for 20% of the trials. During a slow context block of trials, the pursuit target moved at 2 deg/s for 80% of the trials and at 10 deg/s for 20% of the trials. The pursuit target was a patch of 72 dots (half bright and half dark) within a 4 × 4 degree aperture. Both high-contrast (100%) and low-contrast (12-16%) patches of dots were used as targets in equal proportions during these experiments. Every trial started with 800-1600 ms of fixation of a central black spot that was at the center of the patch of dots. After 800-1600 ms, the central spot disappeared and the target underwent 100 ms of coherent local drifting motion before the aperture and dots moved en bloc for an additional 600 ms. On 20% of the trials, a brief, 50-ms pulse of coherent motion at 5 deg/s in the same direction as the pursuit trials occurred after 400 ms of fixation (“early fixation”). On an additional 20% of the trials, the same pulse of motion occurred after 1200 ms of fixation (“late fixation”). The pulse of motion was always followed by smooth pursuit later in the trial. We chose not to present pulse-only trials in order to minimize disruption of motor preparation and expectation. Note that some of the initial speed context data collected in monkey Xt did not deliver any pulses of target motion during fixation.

#### Speed tuning experiments

We assessed the active and passive speed tuning properties of DLPN and NRTP neurons by having the monkeys actively track targets and passively view visual motion at 2, 4, 8, 16, and 32 deg/s. Active pursuit and passive visual motion speed tuning were assessed in separate blocks of trials.

The active pursuit trials used similar stimuli described for the motion pulse experiments with several minor differences. First, the fixation period displayed only a black spot without the patch of dots. Second, after 800-1600 ms of fixation, the patch of dots appeared, the fixation spot disappeared, and 100 ms of coherent local motion began immediately, followed by en bloc movement of the aperture and dots for an additional 600 ms.

During the passive visual motion block of trials, monkeys fixated a red spot that centered on a 16 × 16 deg patch of 288 dots (half were dark and half were light). To prevent the eyes from tracking the moving field of dots and changing the speed of image motion, we imposed a 2 × 2 deg, white central scotoma at the center of the patch of dots. The monkey fixated the stationary stimulus for 800-1600 ms before we delivered 200 ms of coherent local motion of the dots within the 16 × 16 degree aperture. As above, we interleaved randomly trials that presented high- and low-contrast visual motion.

### Quantification

#### Data Analysis

All data analyses were performed using custom code. Eye velocity was computed offline by filtering the horizontal and vertical eye position traces with a zero-phase, two-pole Butterworth low pass filter (cutoff of 25 Hz) and then numerically differentiating the smoothed signals. Saccades were detected automatically using acceleration and velocity thresholds. Both behavioral and neural responses were treated as missing data during saccades. The amplitude of the eye velocity response was taken as the horizontal velocity after rotating the eye velocity vector by the direction of target motion. Time varying firing rates were estimated by smoothing spike trains with a causal post-synaptic current filter (Herzfeld & Beardsley, 2011). Smoothed firing rates then were averaged across single trials of a given condition.

For analyzing preparatory activity, we calculated the average firing rate modulation as the difference between the average firing rate 1400-1500 ms after fixation onset and the average firing rate 150-250 ms after fixation onset. To analyze pursuit-related activity, we calculated the average firing rate during the interval 50-150 ms after the onset of target motion. This interval occurs during the rapid eye acceleration of pursuit initiation and provides an open-loop assessment of responses during the initiation of pursuit before any effects from visual feedback due to imperfect pursuit. We computed neural and eye velocity responses to pulses of visual motion as the maximum minus the minimum firing rate or eye velocity in the interval 0-200 ms after motion pulse onset. In all PSTHs that show response “modulation”, we subtracted the average firing rate in the 100 ms of fixation leading up to motion onset for each neuron’s PSTH before averaging across neurons.

#### Linear model relating the firing rate and eye speed effects of fixation duration and contrast

We computed firing rate modulation across all conditions of contrast, fixation duration, and speed context for each neuron as described above. Then, we normalized firing rates for the range of responses across all stimulus conditions, within each neuron. We then used ridge regression to assign weights relating the normalized activity of each DLPN and NRTP neuron to the average eye speed responses across fixation duration and contrast. We chose to fit the responses for fixation duration/contrast because those were the conditions across our population that showed statistically significant effects. We then used the weights for the best linear model to show that these same neurons could correctly predict the differences in contrast but not the differences across speed context.

#### Linear model relating responses of a model population of MT neurons to the speed tuning response of individual NRTP and DLPN neurons

We generated a model population of area MT that contained 60 units with log-uniformly distributed preferred speeds from 0.2 deg/s to 64 deg/s. Responses (*r*) of a given model MT unit with preferred speed *i* to 2, 4, 8, 16, and 32 deg/s of high contrast visual motion were generated according to:

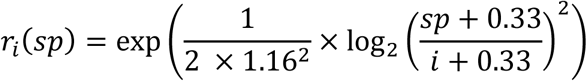

where sp is the visual motion speed input to the model unit. To generate appropriate responses to low-contrast visual motion, we reduced the response amplitudes and shifted the peak towards slower speeds according to:

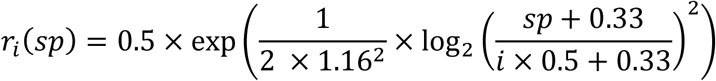

We used population of model MT units as inputs to ask about the properties of a decoder that could account for the responses we recorded each DLPN and NRTP neuron based on inputs from MT. For 5 stimulus speeds and 2 stimulus contrasts, we calculated response modulations to passive visual motion during fixation as the average firing rate 50-250 ms after visual motion onset minus the average firing rate during the 100 ms leading up to motion onset. We normalized the response modulations to the maximum firing rate elicited across all visual motion speeds and contrasts. We then fitted a linear model using regularized ridge regression between the responses of our model area MT population and the normalized responses of each DLPN and NRTP neuron in our sample.

### Statistics

Data were collected from two Rhesus monkeys. For DLPN, we recorded 29 neurons in monkey Xt and 34 neurons in monkey Yo. For NRTP, we recorded 57 neurons each in monkeys Xt and Yo. When we were able to maintain action potential isolation long enough, we ran visual motion pulses and/or speed tuning experiments in multiple directions for a given neuron. For the speed context experiment, we ran 66 experiments on 53 DLPN neurons and 182 experiments on 105 NRTP neurons. For speed context experiments with the added visual motion pulses during fixation, we ran 39 experiments on 29 DLPN neurons and 123 experiments on 93 NRTP neurons. For passive visual motion speed tuning experiments, we ran 106 experiments on 91 NRTP neurons and 66 experiments on 55 DLPN neurons. We included neurons in our sample only if they had at least a 10 spike/s increase in firing rate in at least one condition across speeds and contrast. This left us with 55 experiments from 47 DLPN neurons and 80 experiments from 72 NRTP neurons. Hypothesis testing was performed via two-sided, paired-sample t-tests. We did not formally check our assumption of normality. However, our results were qualitatively and quantitatively unchanged if we used a two-sided Wilcoxon signed-rank test.

## Supplementary information

**Supplementary Figure 1:**
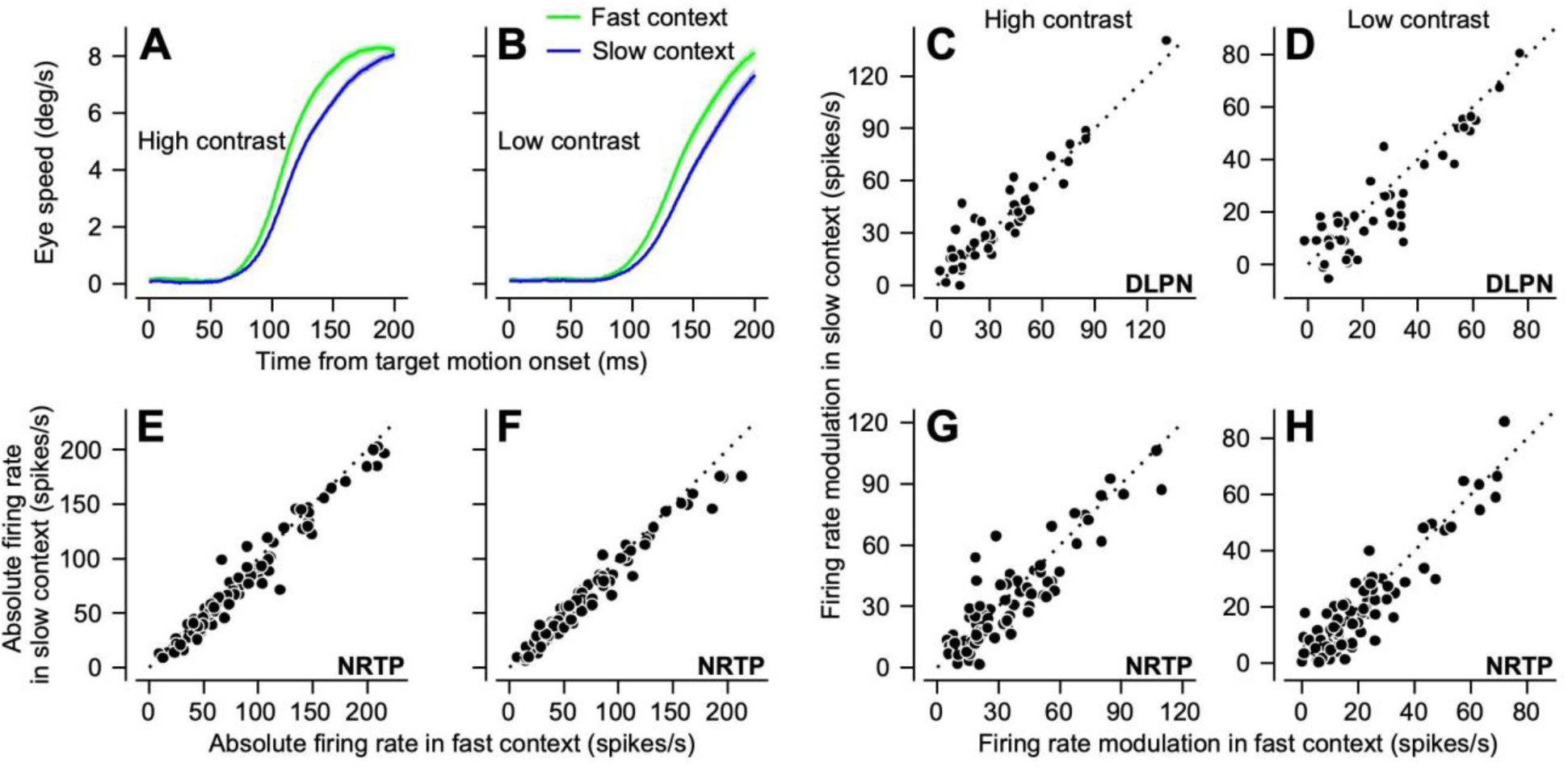
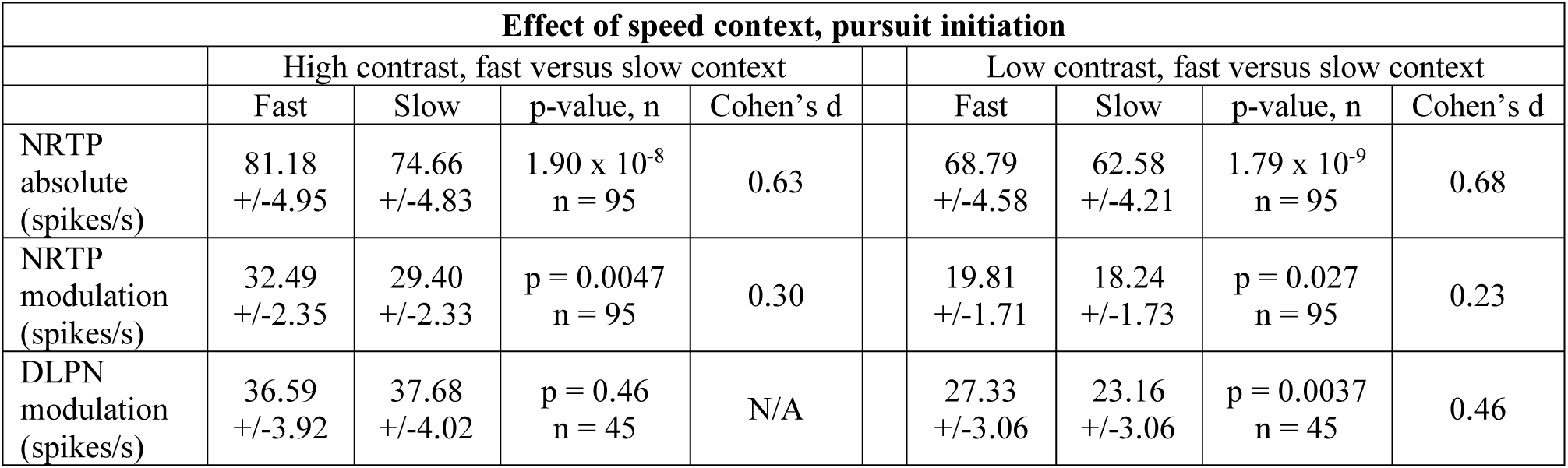
Speed context has small effect on the pontine activity during pursuit initiation. We present data only from 10 deg/s target motions because the visual motion presented in those trials is the same across speed contexts and the only difference is the expectation of target speed according to the speed context it was delivered in. **A, B**: Consistent with previous results, eye speed during the 10 deg/s condition is faster during the fast compared to slow context. **C, D:** In DLPN, pursuit initiation responses in DLPN are affected by speed context for low-contrast, but not high-contrast, targets. **E, F**: The absolute firing rate during pursuit initiation (50-150 ms after onset of target motion) in NRTP in the 10 deg/s probe trials is larger during the fast compared to the slow context for both high- and low-contrast targets, as expected given the higher baseline of preparatory activity. **G, H**: In NRTP, the firing rate modulation during pursuit initiation in the 10 deg/s motion is higher during the fast compared to slow context. Means, standard error of the mean, and hypothesis testing appear in the table, below.

**Supplementary Figure 2:**
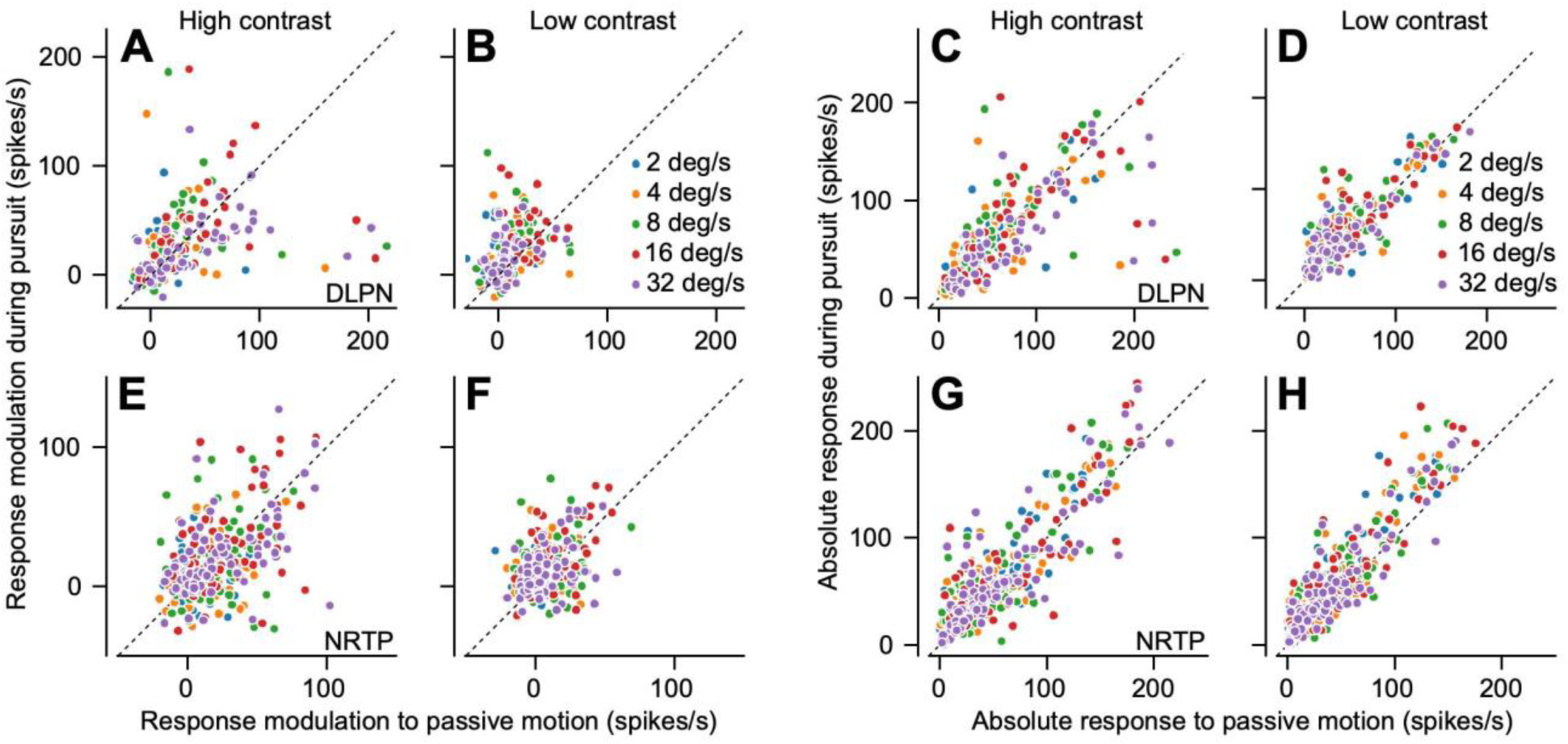
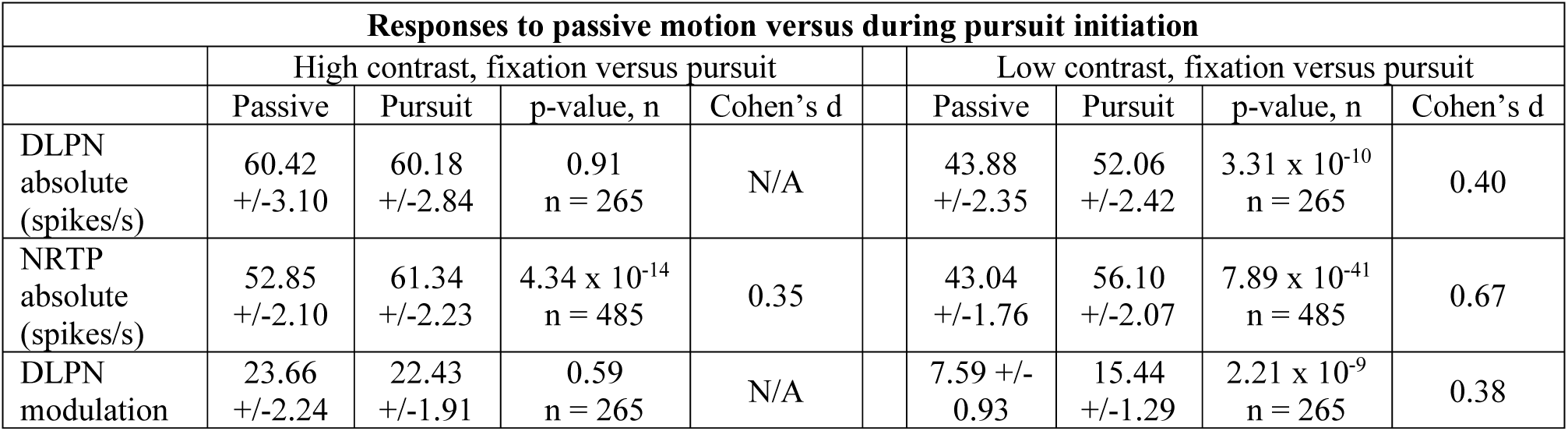
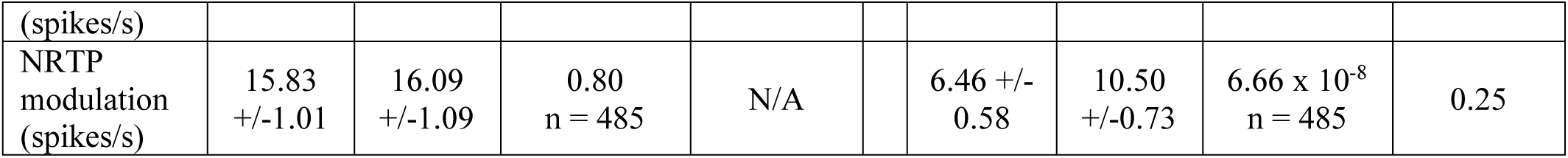
NRTP and DLPN show similar amplitude responses to large field visual motion during fixation versus to active pursuit initiation. Neurons in NRTP and DLPN responded vigorously to visual motion even when it did not drive an eye movement response. Modulation of firing rate is higher for the initiation of pursuit compared to passive visual motion responses only for low-contrast targets (**A, B, E, F**), an effect that is consistent with the idea that extraretinal signals have a larger impact on sensory-motor transformation when sensory input is weak or noisy. Because of the higher baseline provided by preparatory activity, the *absolute* firing rate in NRTP (rather than the *modulation* of firing rate) is larger during active pursuit than for passive visual motion for low- and high-contrast targets (**G, H**). One major caveat that mitigates from drawing strong conclusions from these data is that the visual stimulus differed between the passive visual motion and active pursuit conditions out of necessity. It was a large field with a central scotoma during passive stimulation and was a small patch during pursuit trials (see Methods). **A, B**: each symbol plots data for an individual neuron in DLPN with 5 symbols per neuron for the 5 target speeds. The plot compares response modulation during pursuit versus that for passive visual motion during fixation. The two panels plot data for high-contrast and low-contrast targets separately. **C, D**: Same as **A, B** but plotting absolute firing rate for DLPN. **E, F**: Same as **A, B**, but plotting modulation of firing rate for NRTP. **G, H**: Same as **A, B**, but plotting absolute firing rate for NRTP. Means, standard error of the mean, and hypothesis testing appear in the table, below.

